# Elucidating the conformational dynamics of the mitochondrial localization signal, M3, of TDP-43 and accessing potential inhibitors using molecular docking and simulation

**DOI:** 10.1101/2025.11.13.688389

**Authors:** Ramkumar Balaji, Himanshu Joshi, Basant K Patel

## Abstract

Aberrant mitochondrial localization of the RNA/DNA-binding protein TDP-43 is implicated in amyotrophic lateral sclerosis (ALS) which may affect mitochondrial dynamics and contribute to neuronal toxicity. Inhibitors of the cytoplasmic aggregation of TDP-43 were reported previously, but their effect on the mitochondrial mis-localization of TDP-43 and the mitochondrial function remains to be investigated. Three internal peptide sequences from TDP-43, M1, M3, and M5, were found to enable TDP-43’s mitochondrial localization and competitive inhibition using these peptides thwarted mitochondrial import of TDP-43 and rescued TDP-43-induced cytotoxicity to neurons. Here, we performed a virtual *in silico* screening of 2,115 FDA-approved small molecules for affinity against the M3 region of TDP-43 (aa: 146 – 150) that lies in the RNA recognition motif-1 (RRM1) domain. Multiple all-atom MD simulations were carried out starting with two different conformations of the tandem RRMs to understand the dynamics of the target M3 region in explicit solvent water. The analysis of the simulation trajectories suggests that the M3 region is relatively non-flexible and buried relative to the other regions of the tandem RRM1-2 domains. Cholecalciferol (Vitamin D3), from virtual screening, docked consistently with the M3 region in different docking strategies even amidst the region’s poor solvent accessibility. Vitamin D3 also remained stably bound to the M3 region in most frames of four replica MD simulations, each of one microsecond. Taken together, our study proposes vitamin D3 as a potential binder to the M3 region which may inhibit the pathogenic mitochondrial mis-localization of TDP-43.

## 1.0 Introduction

Neurodegenerative diseases, a significant subset of neurological diseases, are characterized by the death of post-mitotic cells. Under physiological conditions, these cells resist mitotic cues and cell death for longevity and continuous functioning. Prolonged cellular stress causes irreversible injury to the neuronal cells, leading to apoptosis and neurodegeneration. Protein misfolding and aggregation are implicated in dysfunctional proteostasis and significantly contribute to this prolonged cellular stress. Genetic mutations of a specific protein or the mutations of protein folding machinery cause protein misfolding, leading to cellular insults such as mitochondrial dysfunction, inflammation, etc., further leading to increased reactive oxygen species (ROS) and eventually apoptosis (Gandhi et al., 2019). Genes mutate in different brain regions and, based on the cell type affected, lead to different neurodegenerative conditions. Alzheimer’s disease (AD), Parkinson’s disease (PD), Huntington’s disease, and amyotrophic lateral sclerosis (ALS) are the major reported neurodegenerative diseases. Amyotrophic lateral sclerosis is a disease of motor neurons that shows heterogeneity based on the type of motor neurons and regions of the body affected. The clinical manifestation varies from atrophy of the limbs to hyperreflexia, tongue atrophy, or dysfunctional speech. (Taylor et al., 2016). Various proteins and genetic factors are reported to be determinants of ALS, including mutations in the superoxide dismutase (SOD1), TAR-DNA binding protein (TDP-43), fused in sarcoma (FUS), optineurin (OPTN), etc. Among these, TDP-43 is reported to be a significant pathological substrate in ALS and frontotemporal lobar degeneration (FTLD) (Kiernan et al., 2011; Neumann et al., 2006).

TDP-43 is a DNA/RNA binding protein of 414 amino acids organized into three major domains, namely an N-terminal domain, a conserved bipartite RNA-recognition motif (RRM-1 and RRM-2) tandemly placed but separated by a short linker region, and a C-terminal domain with intrinsically disordered regions. These three domains also comprise other physiologically significant areas such as the nuclear localization signal (NLS), nuclear export signal (NES) in the N-terminal and RRM2 domain, respectively, followed by a glutamine/asparagine (Q/N)-rich region, and a glycine-rich region in the C-terminal domain. Internal motifs (M1-M6) for mitochondrial localization, as a stretch of hydrophobic amino acids, are interspersed throughout the length of the TDP-43 (Prasad et al., 2019; Wang et al., 2016).T DP-43, a nuclear resident protein, is reported as ubiquitinated or hyperphosphorylated cytoplasmic inclusions in samples derived from patients with ALS/FTLD (Neumann et al., 2006). The cytoplasmic aggregation of exogenously introduced or overexpressed wild-type (WT) or mutant TDP-43 resulted in enhanced toxicity in various cellular models of ALS (Armakola et al., 2011; Ash et al., 2010; Cascella et al., 2022).

Mitochondrial localization of TDP-43 was reported in the spinal cord motor neurons from ALS patients, and the inhibition of the mitochondrial localization of TDP-43 reduced the toxicity in HEK293 cells. Three mitochondrial motifs M1, M3, and M5 in the N-terminal, RRM1, and C-terminal region respectively were reportedly sufficient for the mitochondrial localization of TDP-43 and the related enhanced toxicity (Wang et al., 2016). However, in the mitochondria, TDP-43 is shown to stabilize the RNA transcripts that code for the electron transport chain (ETC) proteins (Izumikawa et al., 2017). Hence, the context-dependent interaction of TDP-43 with the RNA in the cytoplasm and mitochondria dictates its pathogenicity. Additionally, hyperinflammatory responses associated with ALS are found to be triggered by the release of mitochondrial DNA through the mitochondrial permeability transition pore (mPTP) in HEK293T cells overexpressing TDP-43 (Yu et al., 2020).

Earlier, our group reported that the deletion of a mitochondrial-dependent apoptosis pathway-related gene (*YBH3*), along with cyclin C (*CNC1*) and a mitochondrial fission-related protein (*DNM1*), rescued toxicity in the yeast model of TDP-43 toxicity. (Bharathi et al., 2021). Moreover, toxicity due to a 25 kilodalton fragment of TDP-43 in yeast was mitigated in the presence of an antioxidant, N-acetyl cysteine, suggesting the role of oxidative stress in enhanced toxicity. (Bajpai et al., 2024). These studies suggest that the mitochondrial import of TDP-43 and the oxidative stress associated with the import can also be potential targets for treating the associated neurodegenerative condition.

Small molecules chemically synthesized or biologically derived, along with other biomolecules such as peptides and anti-sense oligonucleotides (ASO), have been widely screened for targeting the aggregation of TDP-43 and other proteins reported in various proteinopathies. Earlier studies from our laboratory have reported small-molecule inhibitors, AIM4 and EGCG, of the aggregation of TDP-43 by biochemical assays and *in silico* techniques. (Girdhar et al., 2020; Meshram et al., 2024). These inhibitors’ effect on the mitochondrial localization of the TDP-43 is yet to be explored, and the efficacy of the inhibitors in the context of the regression of disease in higher animal models is also yet to be established. Hence, screening for inhibitors directly targeting the mitochondrial localization of TDP-43 is also necessary.

Structure-based virtual drug screening is widely used as an alternative to conventional laboratory drug screening. This computational technique reduces the number of drugs tested in the laboratory, reducing cost and effort (Gao et al., 2023). Molecular docking, a method in virtual drug screening, is used to model the interaction between macromolecules and the interaction of small molecules with macromolecules. The mathematical models developed on the physics governing the interatomic interactions lay the foundation of another valuable technique employed in virtual drug screening: Molecular Dynamics (MD) simulation (Hollingsworth and Dror, 2018). MD simulation is employed in sampling the conformations of biomolecules, such as proteins, as they fold or unfold on interaction with other molecules in different environmental conditions (Hollingsworth and Dror, 2018).

Hence, in our current study, virtual screening of 2,115 FDA-approved small molecules is performed against the internal signal sequence M3 for mitochondrial localization in the RRM1 domain of TDP-43. ZINC15, a publicly available database of small molecules, is accessed to obtain the structures of the small molecules to study *in silico*. To understand the dynamics of the target M3 region, an all-atom MD simulation in the presence of explicit solvent is performed on the structure of tandem RRM domains of TDP-43. Two different starting conformations are subjected to three replica one-microsecond simulations each to investigate the dynamics of the M3 region. One is the publicly available solution NMR structure of the tandem RRM domains (PDB ID: 4BS2) without the RNA. The other conformation is the representative structure from clustering the trajectories of the replica simulations of the 4BS2 apo-structure. The M3 region is found to be rigid in all these simulations, as not many fluctuations are seen in the analyzed parameters. Virtual screening against this rigid M3 region with less solvent accessibility was performed on AutoDock Vina, and 246 molecules with a docking score below −8.0 were found. Blind docking was performed with AutoDock 4.2 on these, which reported only four molecules binding at the M3 region. Further screening of the four reported molecules with the modelled full-length TDP-43 reported only Cholecalciferol/ Vitamin D3 (ZINC ID: ZINC4474460) and Atacand (ZINC ID: ZINC3782818), binding to the M3 region. Vitamin D3 is further evaluated for its binding stability with the M3 region by using all-atom MD simulations with multiple starting conformations of the macromolecule.

Vitamin D3 was reported earlier to ameliorate oxidative stress and attenuate neuroinflammation in the mouse models of Parkinson’s disease (PD) and Alzheimer’s disease (AD) (Calvello et al., 2017; Lin et al., 2022; Mehrabadi & Sadr, 2020). However, supplementation of Vitamin D3 orally did not improve motor dysfunction in ALS patients despite the lowered Vitamin D3 serum levels commonly seen in the patients affected, and is probably causing reduced motor functions (Trojsi et al., 2020). Hence, Vitamin D3 can be a potential inhibitor of the mitochondrial localization of TDP-43 and its associated enhanced toxicity.

## 2.0 Materials and Methods

### 2.1 Molecular dynamics simulation of the RRM1-2 apo structure and investigation of the dynamics of the M3 region in the RRM1 domain

All-atom MD simulation was performed using the Groningen Machine for Chemical Simulation (GROMACS), a free software listed under the GNU General Public License. Three replica MD simulations of 1 microsecond each for the RRM1-2 apo-structure (PDB ID: 4BS2 – aa: 102-269) were performed at 310 K in explicit solutions of 150 mM NaCl. The topology of the protein was prepared by the in-built pdb2gmx command using the CHARMM36 forcefield. The complex was solvated in a dodecahedron box using a pre-defined, generic, equilibrated 3-point water solvent model, with the box edges kept 1.5 nm away from the protein. The solvated system was neutralized by adding Na^+^ or Cl^-^ counterions, and additional Na+ and Cl-ions were added to make a 150 mM concentration of NaCl using the genion script of GROMACS. The system finally contained 51,280 atoms, including protein, water, and ions. The whole system was subjected to energy minimization using the steepest descent algorithm to avoid clashes among the atoms in the build configuration, followed by constant number of atoms (N), volume (V), and 300 K temperature (T), *i.e*., the NVT and NPT equilibration for one ns each. Harmonic position restraints to the protein atoms were applied, having the spring constants of 1 kcal mol^−1^ Å^−2^, during the initial 2 ns of equilibration and were later removed during the production run. 1-microsecond production was carried out with a 2-fs integration time step using the NPT ensemble. Periodic boundary conditions and the Particle Mesh Ewald (PME) method over a 12 Å resolution grid were used to calculate the long-range electrostatic interactions (Darden et al., 1993). 310K temperature and 1 atm pressure were maintained using V-rescale and Berendsen coupling methods (Berendsen et al., 1984). Root mean squared deviations (RMSD) for the individual trajectory from three replica simulations were calculated by the built-in “*gmx rms*” command with the first frame from the production run as a reference. Similarly, the individual residues’ root mean squared fluctuations (RMSF) were calculated using the built-in command “gmx rmsf”. In addition, the “*gmx cluster*” built-in command for single linkage clustering of the trajectories was used with the RMSD cut-off of 0.35 nm. A representative structure of each cluster is written in a separate PDB file. The residue-wise percentage relative solvent accessibility (%RSA) and solvent accessible surface area (SASA) were calculated with the Tcl/Tk script in Visual Molecular Dynamics (VMD) (Humphrey et al., 1996). The percentage of relative solvent accessibility of individual residues was calculated using the following formula from the absolute solvent accessibility from VMD.

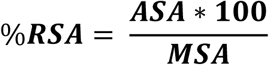

*%RSA – percentage relative solvent accessible surface area for individual residues*

*ASA – Absolute Solvent Accessibility of individual residues measured using VMD in squared Angstrom units (Å^2^) with a 1.4 Å radius probe*.

*MSA – Maximum Solvent Accessibility of the standard amino acid residues reported theoretically earlier in squared Angstrom units (Å^2^) (Tien et al., 2013)*.

The representative structures from the single linkage clustering in Gromacs are color-coded based on the %RSA values using VMD. Additionally, in-house Python 3.10.2 scripts were used to visualize %RSA values for the representative cluster structures as a box plot using Seaborn modules. (Waskom, 2021). The RMSD, RMSF, and SASA plots were generated using GraphPad Prism 10.4.1 (627) software. The above analyses were also performed on the three one-microsecond simulations of the representative structure of the second-largest cluster from the clustering of 4BS2 MD simulation trajectories. The second-largest cluster representative structure is chosen alongside 4BS2 to study the dynamics of M3, as the %RSA of the M3 region in this structure was more than that of the 4BS2 and the rank 1 cluster representative structure.

An R package, Bio3D v2.4-1.9000 (Grant et al., 2021), and Amber’s cpptraj (Case et al., 2023) were used for PCA, considering only the C-α backbone atom of the protein in the trajectories. A ParmEd script converted Gromacs trajectories to the Amber format for Amber PCA analysis. Additionally, hierarchical clustering of the output from PCA was performed using the R package, considering the optimum cluster number as calculated using the Average Silhouette Width (ASW) method. In all the above PCA analyses, the residues at the terminal are neglected as they contribute to most fluctuations in any analysis and contribute to the significant bias. Hence, only residues 106 to 262 are used for rms fit and covariance calculations. The rotational and translational degrees of freedom are removed before the analysis for the entire trajectory using the built-in “*gmx trjconv*” command with relevant arguments to the center and fit options using the starting structure as a reference.

### 2.2 Virtual screening of FDA-approved small molecules against M3

The molecules for virtual screening were retrieved from the ZINC15 (https://zinc15.docking.org). Zinc is a publicly available database of commercially available small molecules. The Tranche Browser of the database allows easy retrieval of the molecules under various categories such as FDA-approved, In-clinical Trials, Metabolites-in-Man, Annotated, etc., (Sterling and Irwin, 2015). The category of molecules chosen in the current study is FDA-approved, and the number of molecules found under this category was 2,115. An NMR structure of the RRM1-2 domain of TDP-43 with the UG-rich RNA bound (PDB ID: 4BS2, amino acids 102-269) (Lukavsky et al., 2013a) was used to screen the molecules against the M3 region. The macromolecular structure was prepared for docking by removing the water molecules, RNA, and the additional sequence from 96 to 101 that does not belong to the TDP-43. The virtual screening was performed in AutoDock Vina v1.1.2, an open-source molecular docking program (Trott and Olson, 2010). A publicly available Perl script for virtual screening was used to screen the molecules. The configuration file carrying the information on the macromolecule and ligand, and the grid box defining the region around M3, was given as input to the Perl script. Based on their dock score, the molecules were categorized as binders to the M3 for further evaluation if their docking score was below −8.0.

Following the defined docking, the prepared macromolecular structure (PDB ID: 4BS2) is blindly docked with the molecules categorized as binders in AutoDock 4.2.6 for further validation of the obtained results. AutoDock 4.2.6 is another general-user docking software requiring grid maps pre-calculated using an autogrid program through the MGL tools v1.5.7 software suite (Morris et al., 2009). The Lamarckian genetic algorithm (GA) was used for the above docking, with the number of GA runs set to 50 in the search parameter.

The top molecules from the two docking strategies were further blindly docked against the modeled full-length apo structure of TDP-43. The modeled full-length TDP-43 apo-structure used in this study is from an all-atom simulation found in the literature after removing the RNA chain (Ingólfsson et al., 2023).

### 2.3 Molecular dynamics simulation for studying the stability of the binding of Vitamin D3 to the M3 region in the apo-protein structure of the RRM1-2 domain

The simulation protocol followed for the protein alone simulations mentioned earlier were repeated for the ligand bound protein complex simulation. The CGenFF server was employed to create the ligand topology and parameter files (Vanommeslaeghe et al., 2012). The topology files of the ligand and protein together constitute the complex topology. The simulation of the ligand-protein complex in 150 mM NaCl solution was carried out with the 4BS2 apo-protein in the rank one representative cluster structure as the macromolecule structure against Vitamin D3. However, four replica simulations of 1 microsecond each were performed for the complex containing the representative structure from cluster two and Vitamin D3, as this structure showed increased solvent-accessibility at M3. The interaction fraction, solvent accessibility of the residues of M3, free energy change for the entire simulation time, and per-residue contribution to the change in free energy were used as metrics to evaluate the stability with which M3 residues bound to the ligand in the simulations. The trajectories and topology files from the protein-ligand simulations were given as input to PyContact, a GUI-based tool for evaluating non-covalent interactions (Scheurer et al., 2018). The tool provides an output of the type of bond and the residue-wise interaction scores for the entire simulation time. This output was saved as a text file, which was used as input for an in-house Python script that calculates the interaction fraction for the residues and type of interaction involved. A value of 1 suggests that hydrogen and other non-covalent interactions are present throughout the simulation. A value of 0.5 for either H-bond/other non-covalent interaction is present for the entire simulation time. As mentioned earlier, the residue-wise SASA for the M3 residues was calculated using VMD. The SASA values were plotted against the M3 residues using GraphPad Prism v10.4.1. The change in free energy and the per-residue contribution to the change in free energy were calculated in the gmx_MMPBSA v1.6.2 tool, an open-source tool to perform end-state free energy calculations for the trajectories from GROMACS simulations (Miller et al., 2012; Valdés-Tresanco et al., 2021).

## 3.0 Results and Discussion

### 3.1 All-atom MD simulations of the RRM1-2 domain in explicit water solvent suggest M3 as a relatively rigid region showing less fluctuation and less solvent accessibility

Each of the tandem RRM domains has a β1-α1-β2-β3-α1-β4-β5 architecture. The internal mitochondrial localization signal, M3, of TDP-43 is in the β3 region of the RRM-1 domain (**Figure S1a**). The three replica simulations of the apo-structure of the tandem RRM domain (PDB ID: 4BS2) varied in their seeding atomic velocities but conformationally had the same starting structure. During the simulations, the apo structure shows deviation from its crystal structure, characterized by root mean squared deviations (RMSD) that saturate around a 12 to 16 Å range after around 700 ns in the 1-microsecond simulations (**Figure 1a**). The single linkage clustering with an RMSD cut-off of 3.5 Å on the concatenated structures from three replica simulations resulted in 399 clusters (**Figure S1b**). The top 3 clusters occupied 17.3%, 14.44%, and 12% of the total submitted structures. The relative solvent accessibility for the M3 residues varied significantly among the clustered structures (**Figure S1c**). Among the various clusters, the representative structure from cluster two was considered for further evaluation along with 4BS2, as the solvent accessibility is higher compared to 4BS2 and the rank one representative structure (**Figure 1b-c**). Unlike the 4BS2 three replica simulations, not all replicas converged around the same RMSD value for the rank two structure. One of the three replicas converged around the RMSD value of 17 Å, whereas the other two converged around 8 Å (**Figure 1d**). However, consistently, in all the protein alone simulations performed, the residues Phe-147 and Phe-149 showed more solvent accessibility than the residues Gly-146, Gly-148, and Val-150 (**Figure 1e-f**). Neither of these starting conformations showed fluctuations in the solvent accessibility of the M3 region. Earlier studies have suggested increased dynamics of the RRM1-2 domain of TDP-43 in the absence of RNA (Donald Scott et al., 2024). The principal component analysis of the 4BS2 trajectory structures in the current study suggested that RRM1 fluctuates more than RRM2 in the explicit solvent. However, the PCA on the concatenated trajectories of the rank 2 cluster showed comparable fluctuations in both domains. Interestingly, dynamics along the essential space could not capture the fluctuations in the M3 (**Figure S2a-f**), suggesting that the beta-sheet (β3) region containing M3, with the surrounding beta-sheets (β1 and β2), resists huge fluctuations in the studied equilibrium MD simulations.

**Figure 1:**
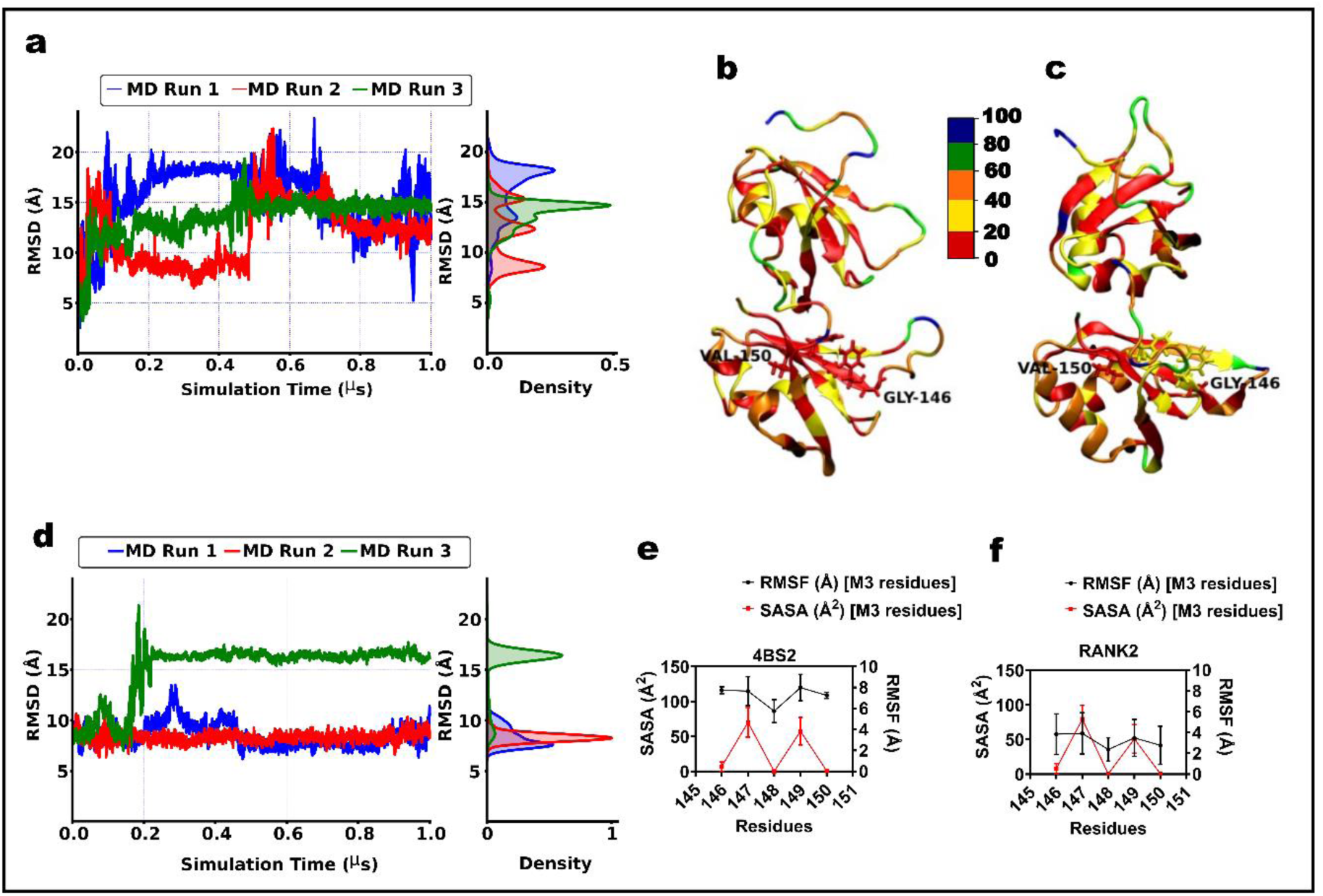
Molecular dynamics simulation of the RRM1-2 apo-structure of TDP-43 with special emphasis on the mitochondrial localization signal, M3. **a)** The RMSD of the backbone C-α of the protein in the three-replica simulations of the 4BS2 apo-structure as a function of simulation time. The probability distribution of the RMSD is shown as a shaded curve on the right side of each RMSD profile. **b-c)** New cartoon representation of the RRM1-2 domain for the NMR structure 4BS2 and the representative structure from cluster two. The residues are colored based on the % RSA as shown in the color map on an RGB scale; red, green, and blue represent low, moderate, and high % RSA. The residues of M3 (aa: 146 – 150) are represented in licorice in addition to the New Cartoon representation. The residues of M3 in the representative structure from cluster two are comparatively more solvent-accessible than those of 4BS2. **d)** RMSD profiles of the C-α backbone atom of the protein in the three replica simulations of the representative structure from cluster two, along with their probability distribution. **e-f)** The twin y-axis plots of RMSF (Å) and SASA (Å^2^) for the M3 residues calculated for the 4BS2 and cluster 2 representative structure trajectories over the simulation time. The residues Phe-147 and Phe-149 are better solvent accessible compared to the almost completely buried Gly-146, Gly-148, and Val-150 in both cases.

### 3.2. In silico drug screening predicts Vitamin D3 as a consistent binder to the internal mitochondrial localization signal, M3 of TDP-43, in different docking strategies

The M3 region consists of the residues Gly-146, Phe-147, Gly-148, Phe-149, and Val-150 in the RRM1 region of the tandem RRM domains (aa: 102-269) of TDP-43 (**Figure S3a**). The docking of the FDA-approved small-molecules against this region manifested 246 molecules with a docking score below −8.0, and the ligand with the lowest score of −9.9 is a molecule with ZINC database id: ZINC35801098 (**Figure S3b**). Blind docking of the 246 binders to the RRM1-2 structure gave ZINC100378061, ZINC538312, ZINC3915154, ZINC3784182, ZINC3782818, ZINC4474460, ZINC13831130, ZINC4474414, ZINC973, ZINC100017856 as top 10 molecules. However, only 4 of these 10 molecules docked to the targeted M3 region. The four molecules are ZINC3782818 (Atacand), ZINC4474460 (Cholecalciferol), ZINC4474414 (Calderol), and ZINC973 (Avobenzone). The docking scores from both strategies and the type of interaction observed for the top 10 molecules are mentioned in **Table 1**, and the docked poses are shown in **Figure 2b, e,** and **Figure S4a-h**. Among the four molecules docked against the modeled full-length apo-structures of TDP-43, only Vitamin D3 (ZINC ID: ZINC4474460) and Atacand (ZINC ID: ZINC3782818) docked at M3 in a few ligand conformations. Though the two molecules showed comparable docking scores, Vitamin D3 was chosen for further analysis as it docked at M3 for most of its conformations and had a better score than Atacand. The results of the blind docking with full-length TDP-43 for the two molecules that stayed at M3 are given in **Table 2, Figure 2c and 2f**. Cholecalciferol/Vitamin D3 (ZINC4474460) is a fat-soluble vitamin stored in the body fat depots and circulates in the bloodstream as a 25-hydroxy derivative [25(OH)D] (Heaney et al., 2009). The residues of M3 interact with Vitamin D3 mainly through hydrophobic interactions, as is evident from the hydrophobic nature of the molecule and the target region.

**Figure 2:**
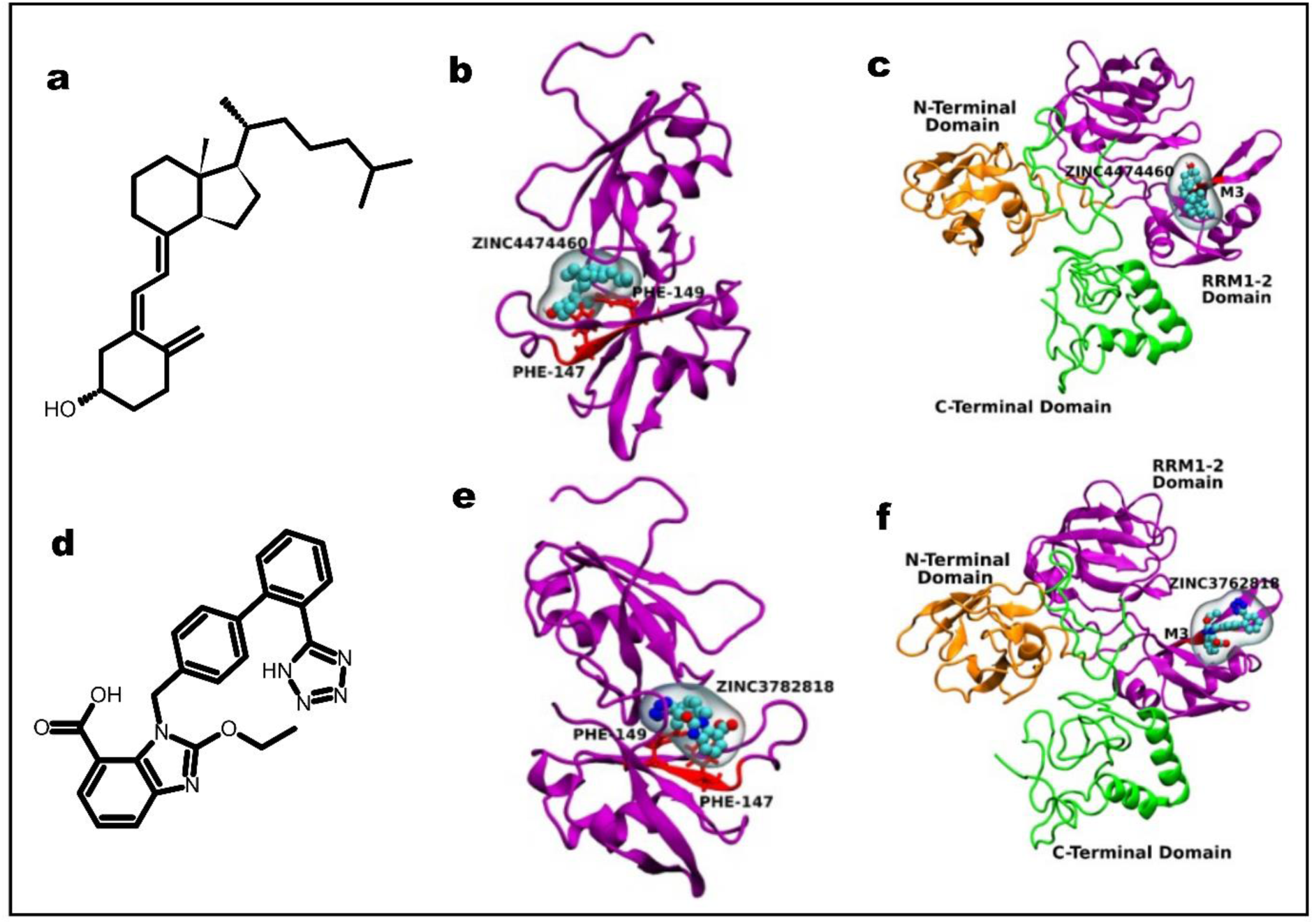
Blind docking poses for the top molecules from defined docking with the solution NMR and the modeled full-length structures of 4BS2 and TDP-43, respectively. **a)** The 2D structure of Vitamin D3 (ZINC ID: ZINC4474460). **b)** Docked pose of Vitamin D3 (ZINC ID: ZINC4474460) with 4BS2 apo-structure. **c)** The docked complex structure of Vitamin D3 with the modelled full-length TDP-43 structures without RNA from the literature (Ingólfsson et al., 2023). **d)** The 2D structure of Atacand (ZINC3782818). **e)** The docked complex structure of Atacand (ZINC ID: ZINC3782818) with the 4BS2 apo-structure. **f)** The docked complex structure of Vitamin D3 with the modelled full-length TDP-43 structures without RNA from the literature (Ingólfsson et al., 2023). The RRM1-2 domain in the above structures is purple in the New Cartoon representation. Van der Waals’ representation of the small molecules is shown along with the MSMS representation. The M3 region is represented as red licorice in addition to the New Cartoon representation.

**Table 1:**
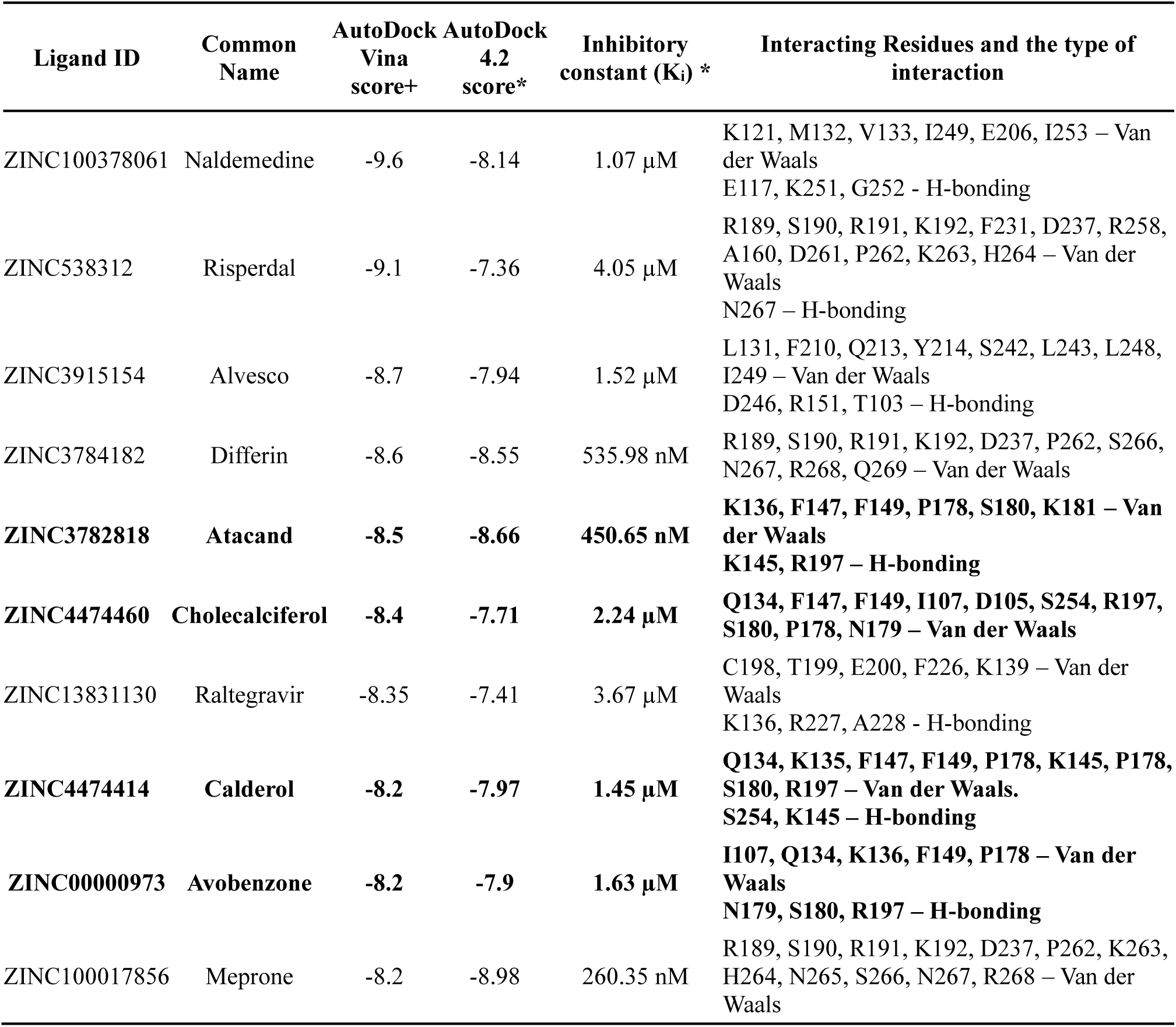
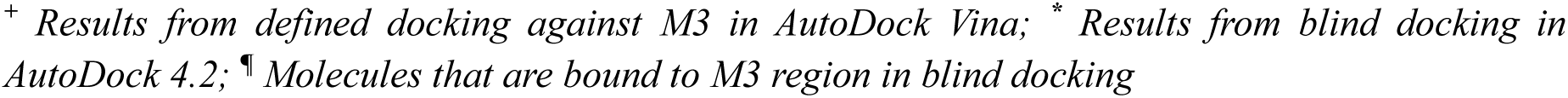
The top molecules ranked based on their defined docking scores against the mitochondrial localization signal M3 in the RRM1-2 domain in AutoDock Vina and blind docking scores against the RRM1-2 domain of TDP-43 in AutoDock 4.2.

| Ligand ID | Common Name | AutoDock Vina score+ | AutoDock 4.2 score* | Inhibitory constant (K<sub>i</sub>) * | Interacting Residues and the type of interaction |
| --- | --- | --- | --- | --- | --- |
| ZINC100378061 | Naldemedine | -9.6 | -8.14 | 1.07 µM | K121, M132, V133, I249, E206, I253 – Van der Waals<br>E117, K251, G252 - H-bonding |
| ZINC538312 | Risperdal | -9.1 | -7.36 | 4.05 µM | R189, S190, R191, K192, F231, D237, R258, A160, D261, P262, K263, H264 – Van der Waals<br>N267 – H-bonding |
| ZINC3915154 | Alvesco | -8.7 | -7.94 | 1.52 µM | L131, F210, Q213, Y214, S242, L243, L248, I249 – Van der Waals<br>D246, R151, T103 – H-bonding |
| ZINC3784182 | Differin | -8.6 | -8.55 | 535.98 nM | R189, S190, R191, K192, D237, P262, S266, N267, R268, Q269 – Van der Waals |
| **ZINC3782818** | **Atacand** | **-8.5** | **-8.66** | **450.65 nM** | **K136, F147, F149, P178, S180, K181 – Van der Waals**<br>**K145, R197 – H-bonding** |
| **ZINC4474460** | **Cholecalciferol** | **-8.4** | **-7.71** | **2.24 µM** | **Q134, F147, F149, I107, D105, S254, R197, S180, P178, N179 – Van der Waals** |
| ZINC13831130 | Raltegravir | -8.35 | -7.41 | 3.67 µM | C198, T199, E200, F226, K139 – Van der Waals<br>K136, R227, A228 - H-bonding |
| **ZINC4474414** | **Calderol** | **-8.2** | **-7.97** | **1.45 µM** | **Q134, K135, F147, F149, P178, K145, P178, S180, R197 – Van der Waals.**<br>**S254, K145 – H-bonding** |
| **ZINC00000973** | **Avobenzone** | **-8.2** | **-7.9** | **1.63 µM** | **I107, Q134, K136, F149, P178 – Van der Waals**<br>**N179, S180, R197 – H-bonding** |
| ZINC100017856 | Meprone | -8.2 | -8.98 | 260.35 nM | R189, S190, R191, K192, D237, P262, K263, H264, N265, S266, N267, R268 – Van der Waals |

<sup>+</sup> Results from defined docking against M3 in AutoDock Vina; <sup>\*</sup> Results from blind docking in AutoDock 4.2; <sup>¶</sup> Molecules that are bound to M3 region in blind docking

**Table 2:** Docking scores of the two FDA-approved small molecules binding at M3 when docked against the modeled full-length TDP-43 structure^#^.

| Ligand ID | Common Name | AutoDock 4.2<br>score<br>(Blind Docking) | Interacting Residues and the type of<br>interaction observed in both<br>structures |
| --- | --- | --- | --- |
| ZINC4474460 | Cholecalciferol | -6.17 | D105, M132, V133, Q134, F149<sup>¶</sup>,<br>K181, R197 – Van der Waals<br>S180 – H-bonding. |
| ZINC3782818 | Atacand | -5.1 | D105, I107, V136, F147<sup>¶</sup>, F149<sup>¶</sup>, S254,<br>I263 – Van der Waals.<br>K145, K136 – Pi-cation Interaction. |

<sup>#</sup> Modelled Full-length TDP-43 structure without RNA, adapted from literature (Ingólfsson et al., 2023); <sup>¶</sup> Residues from the M3 region of TDP-43.

### 3.3. Vitamin D3 stably interacts with the mitochondrial localization signal, M3 of TDP-43, in the four-replica all-atom MD simulations in explicit solvent water

The single-trajectory MD simulation is performed for 4BS2 and the rank-one cluster representative structure docked with Vitamin D3 (ZINC4474460) (**Figure S5a-b**). Four replica MD simulations were carried out for the rank two cluster representative structure with Vitamin D3. The docked complex structure of Vitamin D3 with the rank two cluster representative structure used as a starting conformation for all four replica simulations is shown in **Figure 3a**. The interaction fraction for Vitamin D3 (ZINC4474460) with the rank two cluster representative structure showed that residues Phe-149, Phe-147, and Gly-148 of M3 interact with the ligand for most of the frames from the four replica simulations (**Figure 3b**). The effect of the ligand binding on the solvent accessibility for the M3 residues is insignificant, as the error bars for the SASA calculated for M3 over the entire trajectory for the protein alone and protein-ligand simulations overlap (**Figure 3c**). However, the free energy change for the complex in all four replica simulations is negative (**Figure 3d**). The per-residue decomposition calculated in the gmx MMPBSA tool reported residues Phe-147 and Phe-149 from M3 in two of the four replica simulations of the complex structure, suggesting stable binding of Vitamin D3 with the rank two representative cluster structure of the tandem RRM1-2 domains, and especially near the M3 region (**Table 3**). The principal component analysis of the trajectories of this complex showed similar trends as the protein-alone simulations (**Figure S6a-c**). Hence, the above PCA suggests insignificant differences in the structural conformation of the protein in complex with Vitamin D3 compared with the protein alone simulation for the simulation time considered in this study.

**Figure 3:**
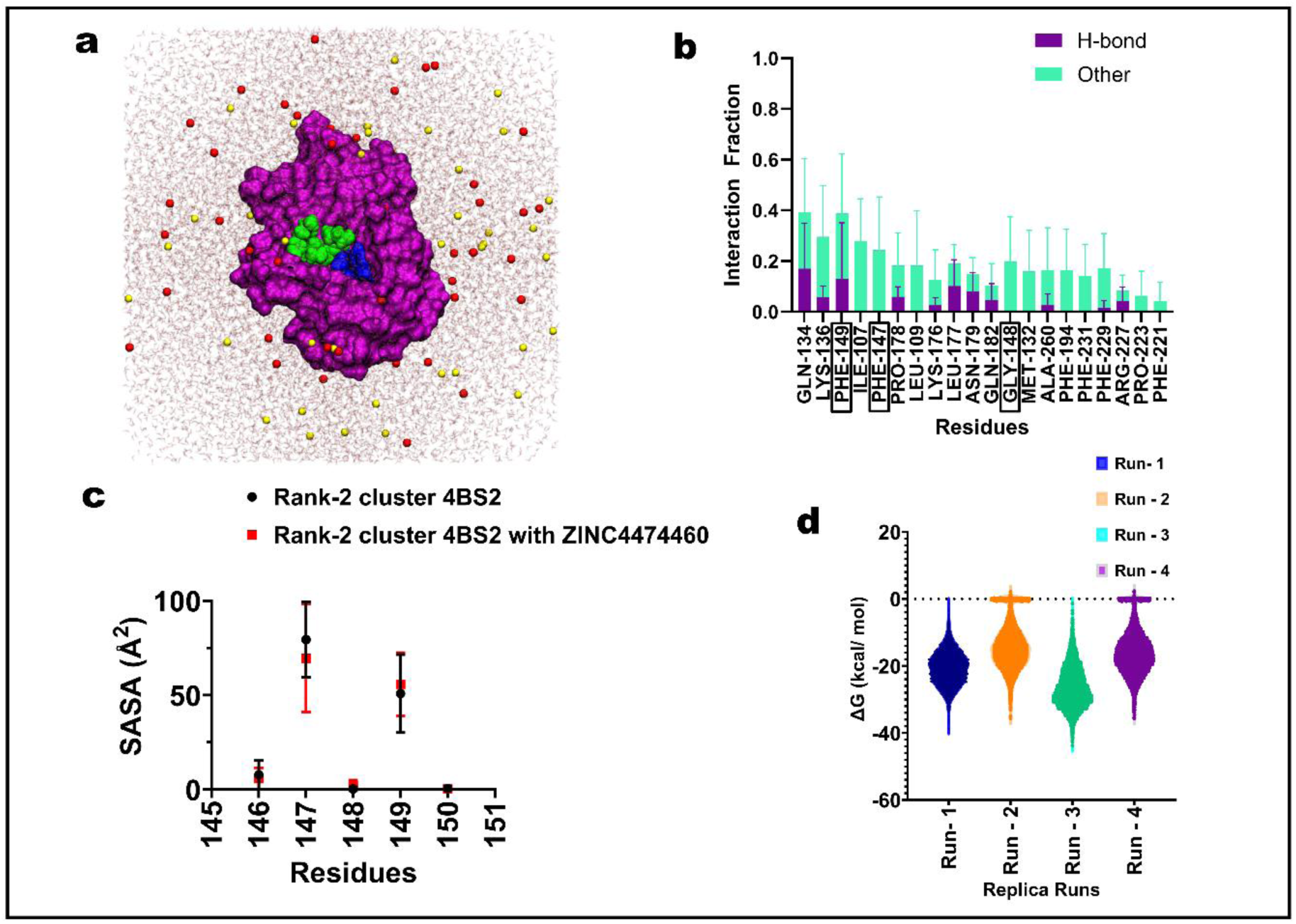
MD simulation results of the rank 2 cluster representative structure with Vitamin D3 (ZINC ID: ZINC4474460) **a)** The all-atom model simulation system of the cluster-2 representative structure of 4BS2 with Vitamin D3. The RRM1-2 domain is shown in the surface representation in purple with the M3 region (aa: 146 – 150) colored in blue. Vitamin D3 is shown in green sphere representation. Line representation of water and sphere representation of ions (red – Na^+^ and yellow – Cl^-^) are shown. **b)** The interaction fraction is represented as a stacked-column chart. The H-bond and other bonds are represented in purple and cyan colors. The error bars represent the standard deviation from the four replica simulations of the rank two structure with Vitamin D3. The residues of M3 (aa: 146 to 150) that are involved in the interaction are highlighted by a rectangular box. The residue Phe-149 interacts with Vitamin D3 in most of the frames. **c)** The Solvent-accessible surface area (SASA) of the M3 residues from the protein alone and protein with Vitamin D3 simulations are shown in black spheres and red squares, respectively. The whiskers represent the standard error from the three and four replica simulations of one microsecond each for the protein alone and the protein with Vitamin D3, respectively. The residues of the M3 showed little difference in their SASA between the protein alone and protein-ligand simulations. **d)** Violin plot of the change in free energy (ΔG) values in kcal/mol for the one-microsecond trajectories of the four replica runs of the rank two cluster representative structure with Vitamin D3.

**Table 3:** Per-residue contribution to the overall change in free energy of binding for the rank 2 cluster representative structure with Vitamin D3.

| Residues | Run – 1* | Run – 2* | Run – 3* | Run – 4* |
| --- | --- | --- | --- | --- |
| ILE-107 | $-1.25 \pm 0.44$ | -- | $-0.56 \pm 0.59$ | -- |
| PHE-147 | $-1.03 \pm 0.48$ | -- | $-0.52 \pm 0.48$ | -- |
| PHE-149 | $-1 \pm 0.58$ | -- | $-1.21 \pm 0.45$ | -- |
| ZINC4474460 | $-11.92 \pm 2.77$ | $-8.96 \pm 4.02$ | $-16.04 \pm 3.48$ | $-9.48 \pm 3.7$ |

\* $\Delta G$ values in kcal/mol ± Standard error values

## 4.0 Conclusion

Aberrant mitochondrial mislocalization of TDP-43 has been reported earlier to increase the toxicity due to TDP-43 in the mammalian cellular models of ALS. The inhibition of this import by peptide inhibitors rescued the cells from the toxicity due to TDP-43, thereby making the mitochondrial import of TDP-43 a potential therapeutic intervention. In this study, virtual screening of the FDA-approved class of small molecules from the ZINC15 database reported Vitamin D3 (ZINC4474460) to be binding at the M3 region in the RRM1-2 domain of TDP-43 with a docking score of −7.71 in the defined docking and −8.4 in the blind docking. Other top-ranked compounds, except Atacand (ZINC3782818), did not bind at the M3 region when blind docking was performed. Moreover, Vitamin D3 bound at the M3 with a docking score of −6.17, when blindly docked to the modelled full-length TDP-43 structure from the literature. Vitamin D3 docked at the targeted M3 region amidst the region showing relatively poor flexibility and solvent accessibility in comparison to the other regions of the RRM1-2 domain, as suggested by the MD simulation analysis of the RRM1-2 domain of TDP-43 in the presence of explicit solvent at 310K. The MM-PBSA analysis of the complex of TDP-43 and Vitamin D3 yielded an average ΔG value (−19.2824 ± 7.58 kcal/mol), with the residues Phe-147 and Phe-149 from the M3 region majorly contributing to the ΔG value in two of the four replica MD simulations of the complex. Hence, Vitamin D3 showed a conspicuous affinity to the M3 region, unlike the other compounds from the tranche, which is further supported by simulations with different starting conformations of the tandem RRM domains of TDP-43. Therefore, Vitamin D3, the fat-soluble vitamin, can be a potential inhibitor to the mitochondrial localization of TDP-43 and its associated toxicity.

## Acknowledgments

We thank IIT Hyderabad, funded by the Ministry of Education (MoE), Govt. of India, for the research infrastructure and support. RB is thankful to MoE, Govt. India for Prime Minister Research Fellowship (PMRF). Himanshu Joshi thanks SERB, Govt. of India, for a research grant (SRG/2022/002109) and DST Inspire Fellowship (IFA20-PH-256). We thank NSM PARAM Seva at IIT Hyderabad for supercomputing time. Basant K Patel thanks SERB, Govt. of India, for a research grant (SERB/CRG/2021/006856).

## Conflicts of Interest

All authors declare no conflicts of interest.

## Notes

### Competing Interest Statement

The authors have declared no competing interest.

