## Supplementary figures for "Elucidating the conformational dynamics of the mitochondrial localization signal, M3, of TDP-43 and accessing potential inhibitors using molecular docking and simulation"

### Supplementary Information

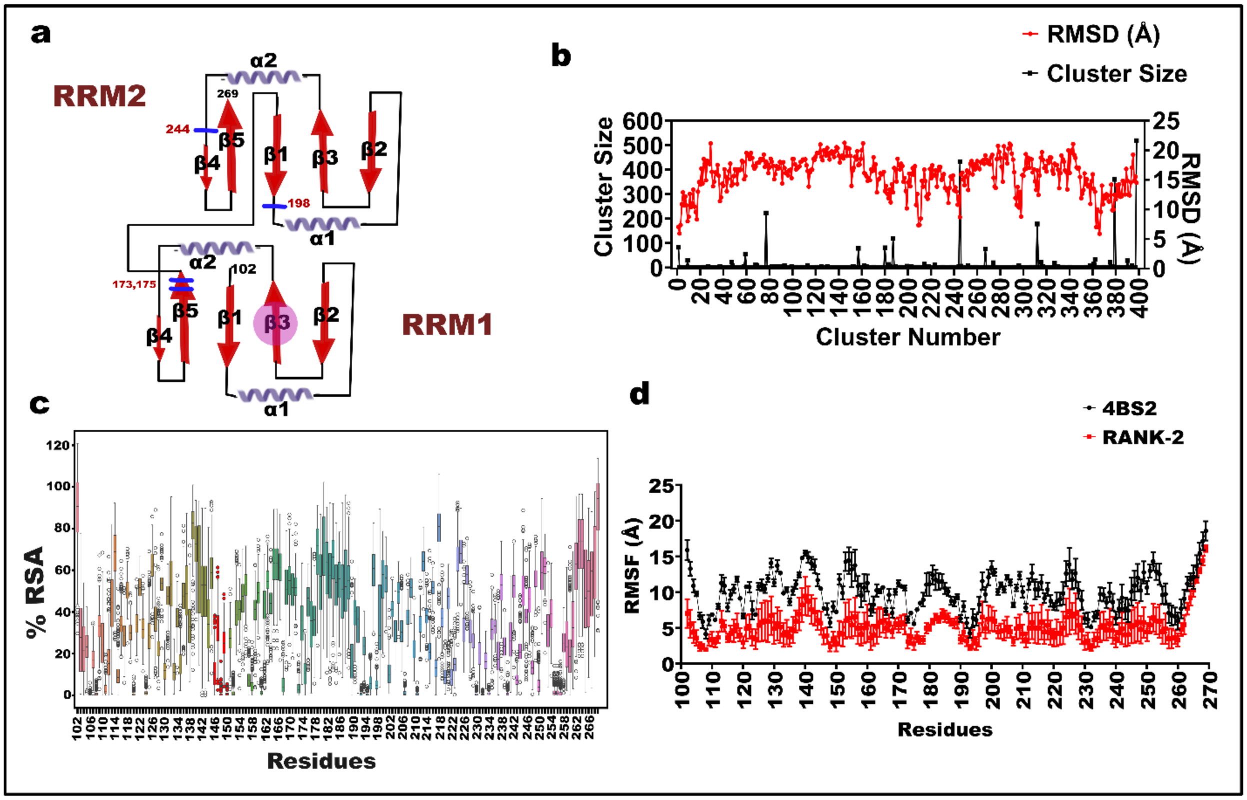


**Figure S1: The single linkage clustering results from the three replica simulation trajectories of the apo-protein 4BS2 structure. a)** The secondary structure organization of the tandem RRM domains of TDP-43. Blue ticks mark the cysteine residues. **b)** The double y-axis plot of the cluster size and RMSD against the cluster numbers. The top cluster (cluster 399) consisted of 519 structures, followed by cluster 246 (rank 2). **c)** The percentage relative solvent accessibility calculated from VMD for the cluster representative structures. Box and whisker plots show the median (middle line), the 75th percentile, and the 25th percentile lining the borders of the box, and the whiskers show the maximum and the minimum values within 1.5 times the Interquartile range (IQR). The additional dots are the outliers that fall outside the range of the whiskers. The red-colored boxes from residue 146 to 150 correspond to the M3 region in the RRM1 domain. The residues Phe-147 and Phe-149 showed more fluctuations, whereas Gly-146, Gly-148, and Val-150 stayed buried consistently. **d)** The Root Mean Square Fluctuations of the residues in the 4BS2 and the rank two cluster representative structure simulations. The 4BS2 structure showed more fluctuations than the rank two cluster representative structure. However, the residues in and around the M3 region showed little fluctuations. The box or the dot represents the mean value, while the whiskers represent the standard deviation from the three replica simulations in each case.

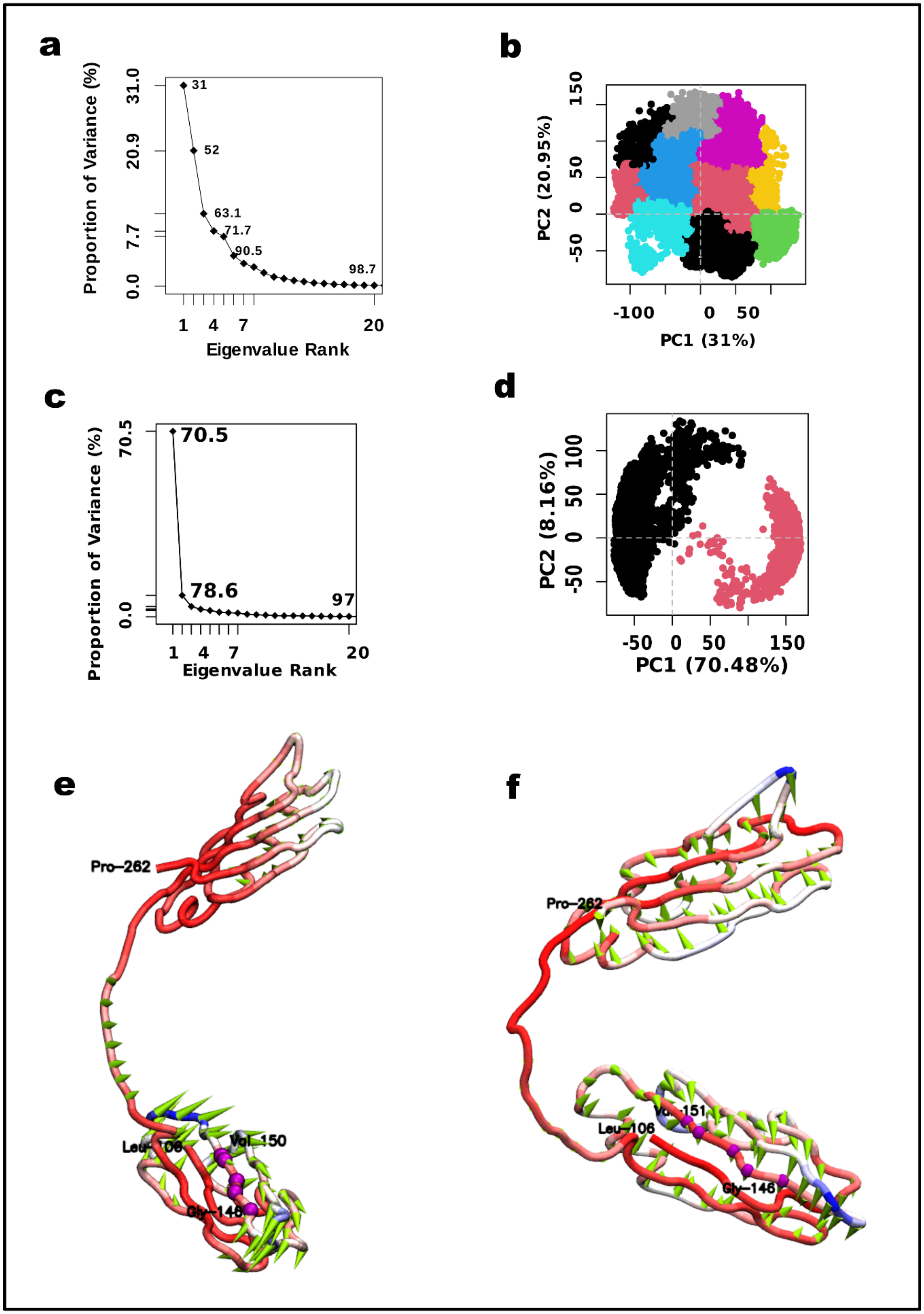

a
Proportion of Variance (%)
31.0
20.9
7.7
0.0
31
52
63.1
71.7
90.5
98.7
1
4
7
20
Eigenvalue Rank
b
PC2 (20.95%)
150
50
0
-50
-100
0
50
PC1 (31%)
c
Proportion of Variance (%)
70.5
0.0
70.5
78.6
97
1
4
7
20
Eigenvalue Rank
d
PC2 (8.16%)
100
50
0
-50
-50
0
50
150
PC1 (70.48%)
e
Pro-262
Val-150
Gly-146
f
Pro-262
Leu-106
Gly-146

**Figure S2: Principal component analysis of simulation trajectories differing by the tandem RRM domain starting conformations to understand the M3 dynamics. a)** The Scree plot shows the proportion of variance against the eigenvectors. The numbers in the plot are the cumulative variance (in percent) for each eigenvector. The first three eigenvectors picked around 60 percent variance and hence are sufficient to capture the concerted motion of the 4BS2. **b)** 2D score plot for the submitted trajectories along the first two principal component axes. The average silhouette width (data not shown) suggested 10 clusters in hierarchical clustering of the PCA scores, and hence, the 2D plot is colored based on their corresponding clusters. **c)** Scree plot for PCA on the three replica simulation trajectories of the rank two cluster representative structure, showing significant variations picked by the first principal component axis alone. **d)** Hierarchical clustering into optimum clusters suggested by the Average Silhouette Width (ASW) is two for the PCA analysis of trajectories from three replica simulations of the rank two cluster representative structure, as shown in two colors in the 2D PCA score plot. **e)** Porcupine plot of the backbone C-α atoms of the 4BS2 simulation structures used for PCA analysis. The spikes represent the axis along which the movement occurs, and the spike's length represents the motion's amplitude. The tube representation of the molecule is used, and the color code represents the mobility of the residues, with red being the least mobile while blue suggests more mobility. RRM1 seems more mobile than the RRM2 domain, but the residues of the M3, represented as purple Van Der Waals spheres, seem less mobile. **f)** Porcupine plot for the PCA analysis on the rank two cluster representative structure simulation trajectories. The fluctuations of the M3 region are less like the 4BS2 simulation; however, the RRM2 domain showed more fluctuations than in the 4BS2 structure.

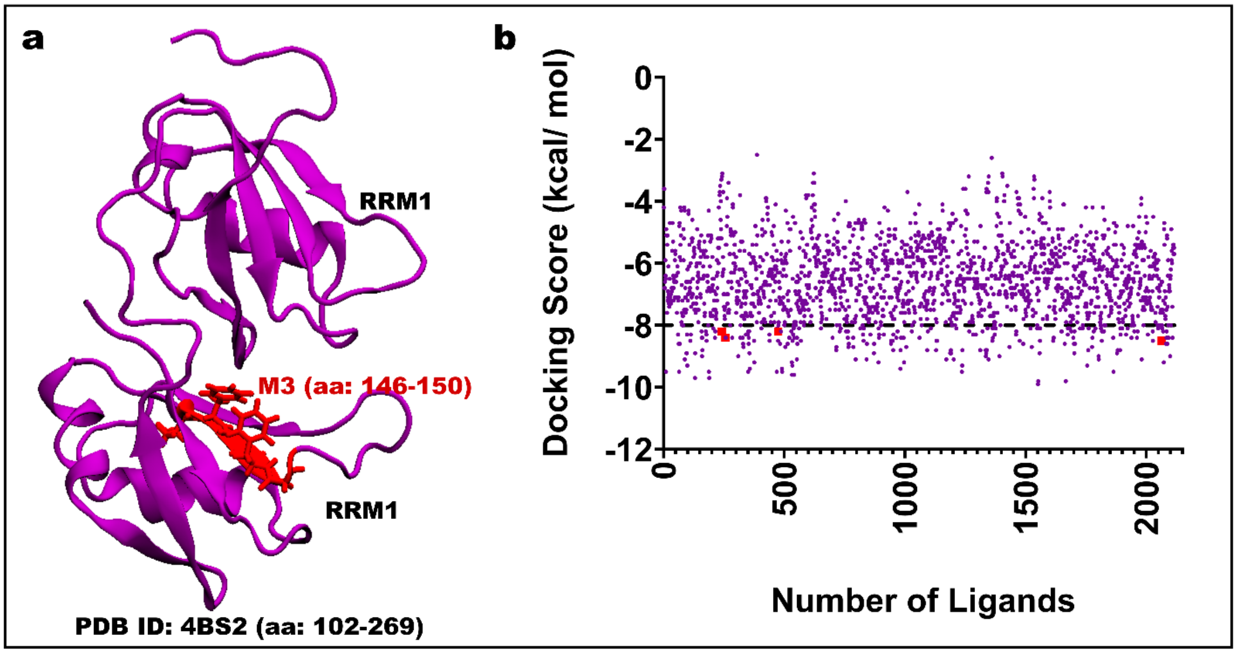


**Figure S3 - The NMR structure of the tandem RRM domains of TDP-43 and scatter plot of the results of the virtual screening of FDA-approved compounds from the ZINC database against M3: a)** The NMR apo-structure of the tandem RRM domains of TDP-43 (RCSB PDB: 4BS2) with their linker region in New Cartoon Representation. The internal mitochondrial motif M3 in the RRM1 domain is colored red, while the remaining domain regions are purple. **b)** Defined docking results of the FDA-approved small molecules from the ZINC library against the M3 region of the TDP-43. The four compounds interacting with M3 in the blind docking are red squares, and the remaining small molecules are purple circles in the scatter plot. The axis for the docking score -8.0 is represented as a dashed line since a docking score < -8.0 is considered a separation criterion for binders and non-binders in the study.

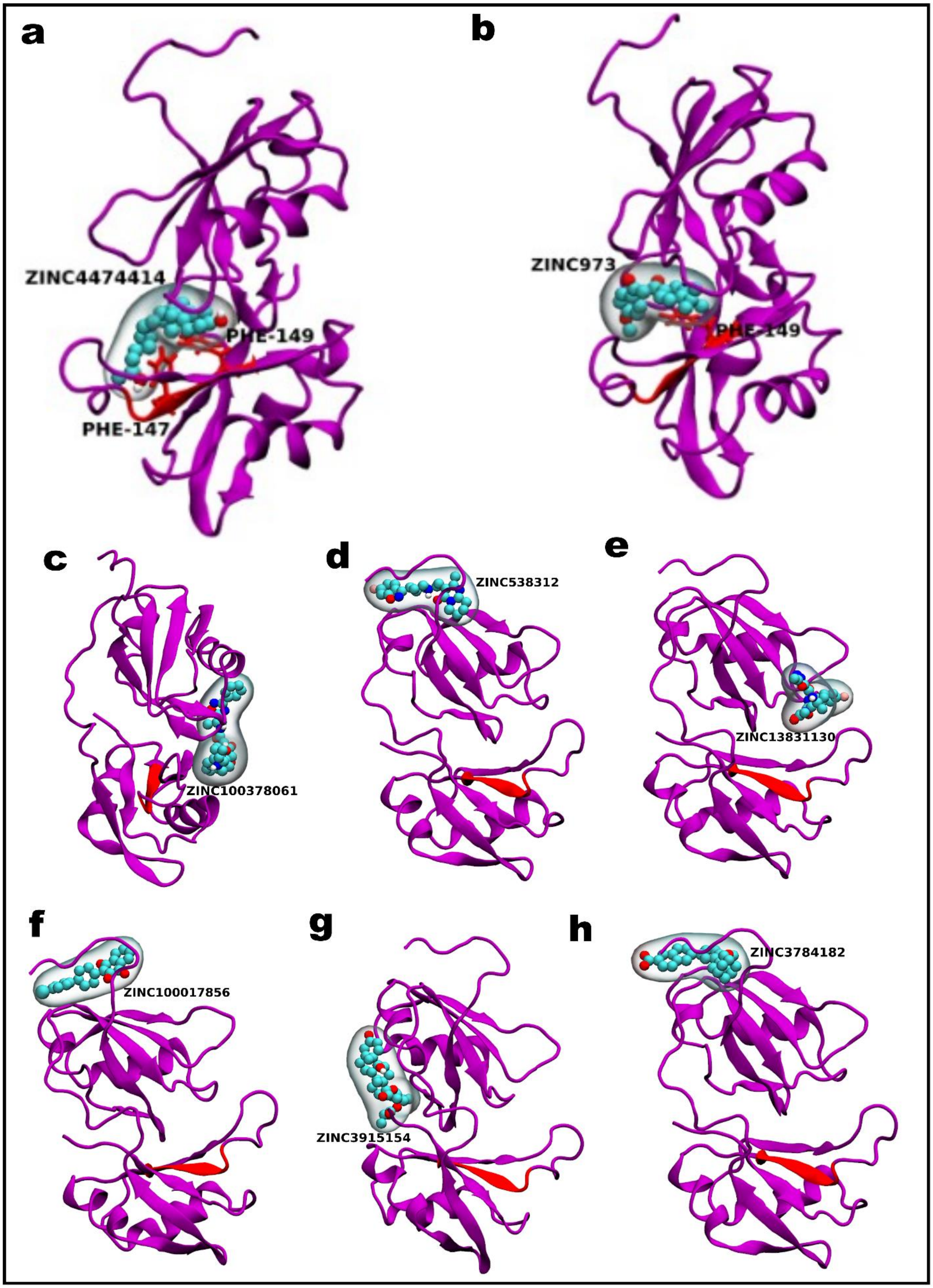
a
ZINC4474414
PHE-149
PHE-147
b
ZINC973
PHE-149
c
ZINC100378061
d
ZINC538312
e
ZINC13831130
f
ZINC100017856
g
ZINC3915154
h
ZINC3784182

**Figure S4: Blind docking poses for the top-scored ligands with the tandem RRM domains of TDP-43. (a-b)** Docking poses from blind docking for the ligands Calderol (ZINC ID: ZINC4474414) and Alvesco (ZINC ID: ZINC973) that docked at M3 in the RRM1 domain of TDP-43. **(c-h)** The docked poses from blind docking of the ligands Naldemedine (ZINC ID: ZINC100378061), Risperdal (ZINC ID: ZINC538312), Raltegravir (ZINC13831130), Meprone (ZINC100017856), Alvesco (ZINC ID: ZINC3915154), and Differin (ZINC ID: ZINC3784182), that gave the top scores in the blind and defined docking with the apo-protein structure of TDP-43 (PDB ID: 4BS2). These compounds did not bind to the M3 region at the blind docking, though their docking scores were below -8.0 kcal/ mol. A purple New Cartoon representation is shown for the tandem RRM1-2 domain, with the M3 region alone colored in red. The small molecules are shown in the Van Der Waals and blown-glass representations using VMD.

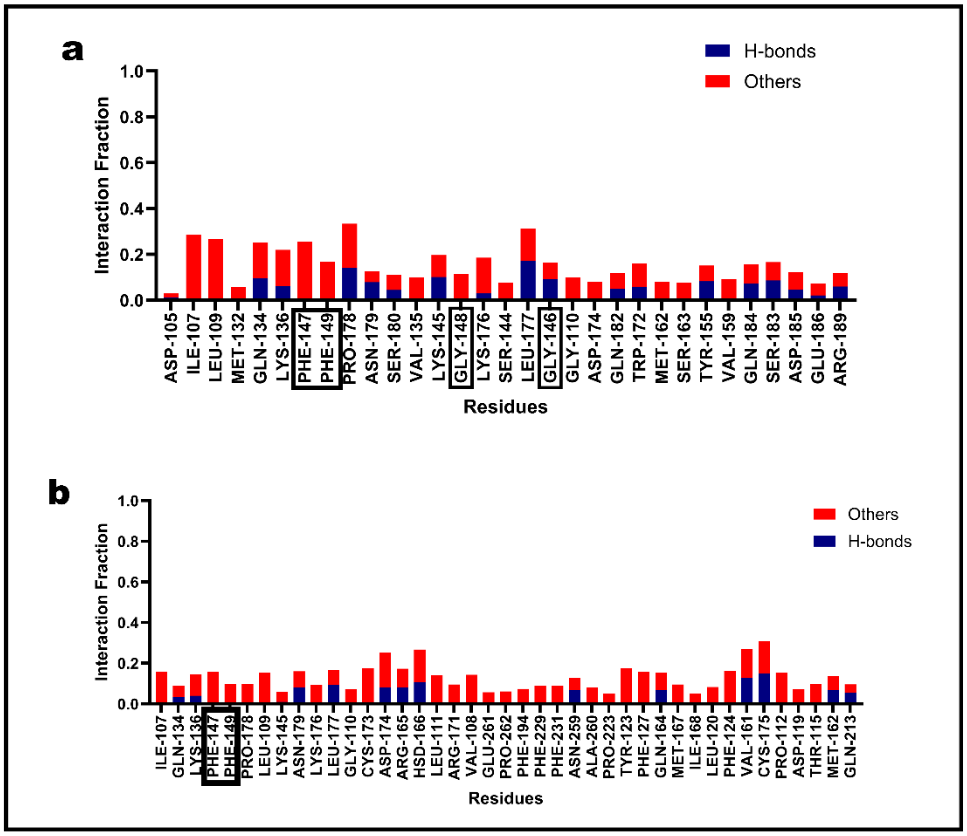


**Figure S5: Interaction fraction profile from the simulation of the 4BS2 structure and Rank–1 representative cluster structure with Vitamin D3 (ZINC ID: ZINC4474460). a)** The interaction fraction from the one-microsecond simulation of Vitamin D3 with the 4BS2 structure.

**b)** The interaction fraction from a one-microsecond simulation of Vitamin D3 with the rank one cluster representative structure from clustering of the three replica 4BS2 simulations. The residues of the M3 are marked by a box in both cases. ZINC4474460 (Vitamin D3) interacted with the M3 region in most frames of the 1-microsecond simulation in the presence of explicit water solvent.

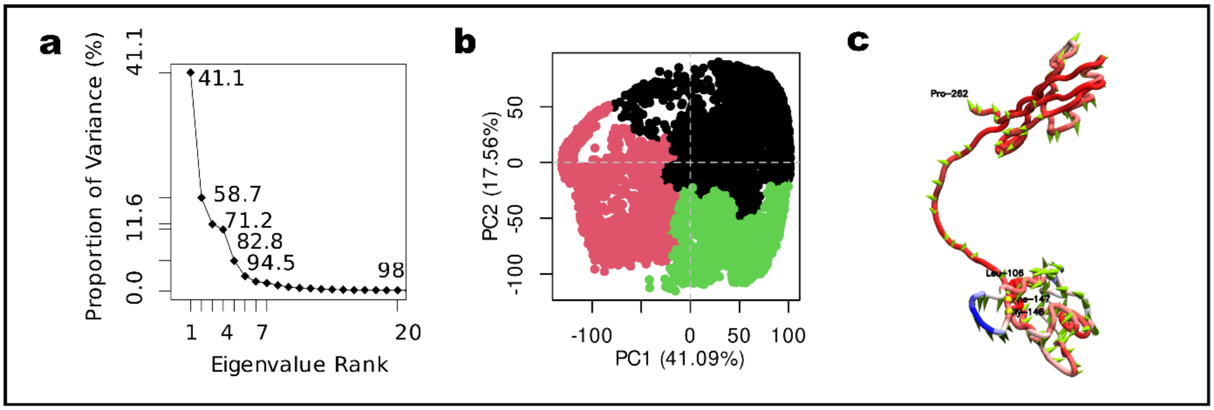


**Figure S6 – Principal component analysis of the trajectories from the four 1-microsecond simulations of the rank two tandem RRM domain structure with Vitamin D3 (ZINC ID: ZINC4474460). a)** The scree plot showing the proportion of variance along the eigenvectors from the PCA analysis performed on the protein C-α backbone atom of the rank two protein structure from the four replica rank 2-ZINC4474460 simulations. The first three vectors, forming the principal component axes, captured almost 70 percent of the variance. **b)** The ASW method suggested three as the optimum clusters formed by the hierarchical clustering represented in the three colors in the 2D PCR score plot for the input trajectories. **c)** Porcupine plot of the protein C-α backbone atom for the rank two cluster representative structure after PCA analysis from the rank 2 cluster representative structure– ZINC4474460 simulations.

**Table S1:** The components of the free energy calculations and their contributions as calculated by gmx_MMPBSA for the Rank 2 cluster representative structure with Vitamin D3 four-replica simulations of one microsecond each.

| Energy Components | RANK 2 cluster representative structure of 4BS2 + Vitamin D3 | | | |
|---|---|---|---|---|
| | **Run - 1¶** | **Run - 2¶** | **Run - 3¶** | **Run - 4¶** |
| VDWAALS ($E_{VdW}$) | -27.88 ± 0.06 | -20.9 ± 0.09 | -36.6 ± 0.07 | -21.76 ± 0.08 |
| EEL ($E_{ele}$) | -2.61 ± 0.04 | -2.24 ± 0.04 | -1.16 ± 0.04 | -1.98 ± 0.04 |
| EPB ($E_{polar}$) | 13.31 ± 0.06 | 11.69 ± 0.07 | 15.45 ± 0.05 | 10.65 ± 0.06 |
| ENPOLAR ($E_{nonpolar}$) | -3.49 ± 0.01 | -2.72 ± 0.01 | -4.16 ± 0.01 | -2.75 ± 0.01 |
| GGAS ($E_{VdW} + E_{ele}$) | -30.49 ± 0.08 | -23.13 ± 0.11 | -37.76 ± 0.08 | -23.73 ± 0.1 |
| GSOLV ($E_{polar} + E_{nonpolar}$) | 9.82 ± 0.05 | 8.97 ± 0.06 | 11.3 ± 0.05 | 7.9 ± 0.05 |
| **Total ($E_{VdW} + E_{ele} + E_{polar} + E_{nonpolar}$)** | **-20.67 ± 0.05** | **-14.16 ± 0.07** | **-26.46 ± 0.06** | **-15.84 ± 0.06** |

¶The values are in kcal/mol ± Standard error of the means (SEM) over the trajectory frames used.
